# Computational performance and accuracy of Sentieon DNASeq variant calling workflow

**DOI:** 10.1101/396325

**Authors:** Katherine I. Kendig, Saurabh Baheti, Matthew A. Bockol, Travis M. Drucker, Steven N. Hart, Jacob R. Heldenbrand, Mikel Hernaez, Matthew E. Hudson, Michael T. Kalmbach, Eric W. Klee, Nathan R. Mattson, Christian A. Ross, Morgan Taschuk, Eric D. Wieben, Mathieu Wiepert, Derek E. Wildman, Liudmila S. Mainzer

**Affiliations:** National Center for Supercomputing Applications, University of Illinois at Urbana-Champaign, USA; Mayo Clinic, Department of Research Services, Rochester, MN, USA; Mayo Clinic, Department of IT Executive Administration, Rochester, MN, USA; Mayo Clinic, Department of Health Sciences Research, Rochester, MN, USA; Institute for Genomic Biology, University of Illinois at Urbana-Champaign, USA; Department of Crop Sciences, University of Illinois at Urbana-Champaign, USA; Mayo Clinic, Department of Biochemistry and Molecular Biology, Rochester, MN, USA; Department of Molecular and Integrative Physiology, University of Illinois at Urbana-Champaign, USA; Genome Sequence Informatics, Ontario Institute for Cancer Research, Toronto, Ontario, Canada

**Keywords:** Sentieon, benchmarking, variant calling

## Abstract

As reliable, efficient genome sequencing becomes more ubiquitous, the need for similarly reliable and efficient variant calling becomes increasingly important. The Genome Analysis Toolkit (GATK), maintained by the Broad Institute, is currently the widely accepted standard for variant calling software. However, alternative solutions may provide faster variant calling without sacrificing accuracy. One such alternative is Sentieon DNASeq, a toolkit analogous to GATK but built on a highly optimized backend. We evaluated the DNASeq single-sample variant calling pipeline in comparison to that of GATK. Our results confirm the near-identical accuracy of the two software packages, showcase perfect scalability and great speed from Sentieon, and describe computational performance considerations for the deployment of Sentieon DNASeq.

## 1. Introduction

Advancements in sequencing technology [1, 2] have resulted in an explosion of Whole Genome Sequencing (WGS) and Whole Exome Sequencing (WES) [3]. As sequencing machines become faster and cheaper [4], analysis must speed up as well. It is no longer acceptable for genomic variant calling to take days or even hours on a single WGS sample. Yet the standard community-accepted software package GATK (Genome Analysis Toolkit [5]) still requires the use of multiple nodes for many hours, even after optimization [6].

Since 2014, Sentieon’s DNASeq pipeline [7] has offered an appealing alternative to GATK. Other ultrafast software alternatives, such as Genalice [8] and Isaac [9], do not adhere to the original algorithms and file formats of GATK. However, Sentieon follows the GATK Best Practices [10, 11] and reimplements the same algorithms in C, C++, Python and ASSEMBLY. It thus boasts a highly optimized rewrite of the Java-based GATK, which helps facilitate its adoption in research and the clinic. It also includes an optimized version of the popular BWA MEM aligner [12].

This whitepaper presents the results of unbiased bench-marking by an independent academic group. It confirms Sentieon’s impressive compute speed, accomplished without loss of accuracy in comparison to GATK3.8 and GATK4. The work is focused on the broadly applicable and clinically useful case of single-sample variant calling.

## 2. Methods and Results

### 2.1 Experimental setup

#### Software versions

All Sentieon tests were run using version 201711.02, except for one BWA MEM performance benchmark run on version 201711.03. A trial license was provided by Sentieon. GATK3.8 was downloaded from the Broad Institute’s software download page [13], build GATK-3.8-0-ge9d806836. Picard version 2.17.4 and GATK 4.0.1.2 were downloaded from GitHub as pre-compiled jar files.

#### Tools Benchmarked

We benchmarked “best practices” single-sample variant calling pipelines built with GATK3.8, GATK4.0 and Sentieon DNASeq (Table 1, see section 2.2 for exceptions). Alignment was performed by BWA or its improved alternative from Sentieon (marked with †).

**Table 1.** Sentieon DNASeq vs. GATK pipelines.

| Pipeline Step | Sentieon | GATK 3.8/4.0 |
| --- | --- | --- |
| Alignment | BWA MEM† | BWA MEM |
| Sorting | Sort utility | NovoSort |
| Deduplication | LocusCollector and Dedup | MarkDuplicates |
| Realignment | Realigner | *Not recommended* |
| Quality score recalibration | QualCal | BaseRecalibrator |
| Apply new quality scores | *Invoked during Haplotype* | PrintReads (3.8)/ ApplyBQSR (4.0) |
| Variant Calling | Haplotype | HaplotypeCaller |

#### Data

Three datasets were used in testing: (1) A dataset corresponding to whole genome sequencing (WGS) of NA12878 [14, 15] to ∼20X depth was downloaded from Illumina BaseSpace on Dec 16, 2016. The NA12878 Illumina Platinum variant calls were used as the truth set to assess variant calling accuracy on these reads. (2) A dataset corresponding to WGS of NA24694 [16] was downloaded on January 19, 2018 from Genome in a Bottle (GIAB) [17]. The NA24694 data arrived in multiple files, which we merged into several subsets to mimic sequencing depths of 25X, 50X, 75X, and 100X. (3) A small synthetic dataset simulating WGS on chromosomes 20-22 was created using NEAT-genReads [18, 19], a synthetic reads generator. The software introduced random mutations into the hg38 reference, and the simulated reads were produced from that mutated reference. The mutations were recorded in the “Golden VCF”, which was used as the truth set when assessing variant calling accuracy on these synthetic data. Command used to generate synthetic reads and Golden VCF:

~~~
python genreads.py -r hg38_chr20_21_22.fa \
   -R 100 --pe 300 30 \
   -M 0.001 -E 0.001 \
   --bam –vcf
~~~

The October 2017 GATK bundle was used for the human reference (hg38), dbSNP (build 138), and the Mills and 1000G gold standard indels.

#### Hardware

All tests were conducted on Skylake Xeon Gold 6148 processors with 40 cores, 2.40 GHz. Each node had 192 GB, 2666 MHz RAM. The nodes were stateless, connected to a network-attached IBM GPFS ver. 4.2.1 with custom metadata acceleration. The cluster used EDR InfiniBand with 100 Gb/sec bandwidth, 100 ns latency. Nodes ran Red Hat Enterprise Linux 6.9. Each test was run on a single node. We ran 2-3 replicates of most tests; the difference across the replicates was negligible, and we are confident that the walltime was not affected by other activity on the cluster.

### 2.2. Tool comparison overview: Sentieon vs GATK

Sentieon DNASeq tools largely mirror those of the GATK (Table 1), and in both toolkits the steps can be individually swapped in and out of a pipeline or replaced by other tools. However, although GATK no longer recommends realignment, that step can convey benefits in a Sentieon pipeline. Thus we included Sentieon’s Realigner tool in all runs except for those intended to compare directly to GATK. Another difference is that GATK creates a separate recalibrated BAM by default, using PrintReads or ApplyBQSR, depending on version. However, Sentieon’s default is to apply BQSR calculations “on the fly” during the Haplotyper step without generating a separate recalibrated BAM, which results in performance improvement by reducing I/O to disk and avoiding a proliferation of intermediary files. Sentieon does have the option to generate a recalibrated BAM with its ReadWriter algorithm, but there is no need for an equivalent of the PrintReads/ApplyBQSR step of GATK in the DNASeq pipeline.

### 2.3. Variant calling accuracy

The winning accuracy of Sentieon’s DNASeq pipeline has already been demonstrated in several FDA and DREAM Challenges [20, 21, 22]. Nevertheless, we ran a cursory comparison of Sentieon’s DNASeq accuracy against the newly released GATK4. The NA12878 and the synthetic chr 20-22 datasets were run through both pipelines. The resultant VCFs were compared to the respective truth sets and to each other, using the vcf-compare tool from the NEAT package [23]. The comparison was limited to the Illumina Platinum confident regions [24]. Command used to run comparisons:

~~~
python vcf-compare.py -r hg38.fa
   -g golden_truth.vcf -w workflow.vcf \
   --vcf-out -o output_directory \
   -t ConfidentRegions.bed -T 0 \
   --incl-homs --incl-fail --no-plot
~~~

Concordance was defined as the percentage of variants present in the truth sets that were correctly identified by the respective softwares. In comparing Sentieon and GATK4 directly, we treated the output from GATK4 as the truth set. Sentieon and GATK4 were highly concordant with each other (Table 2) on both datasets, as expected due to their nearly identical algorithms. Using realignment in DNASeq slightly improved the concordance to GATK4, and the difference could become more significant on datasets of poorer quality. Both toolkits had high rates of variant detection relative to the truth sets. Significantly, GATK and Sentieon demonstrated identical detection rates on the Illumina Platinum data.

**Table 2.** Variant detection accuracy in Sentieon DNASeq and GATK 4.

| Dataset: | Synthetic WGS, chr 20-22 | NA12878 |
| --- | --- | --- |
| Sentieon vs. GATK 4 | 99.91% | 99.78% w/ Realigner<br>99.72% w/o Realigner |
| Truth sets: | Golden VCF | Illumina Platinum variant calls |
| Sentieon vs. Truth set | 92.80% | 94.9% |
| GATK4 vs. Truth set | 91.14% | 94.9% |

### 2.4. Thread-level scalability

We tested the single-node scalability of Sentieon DNASeq by running the same pipeline with 4, 8, 16, 24, and 40 (max) threads. All the constituent tools appeared to scale equally well (data not shown), and the entire DNASeq pipeline scaled near-perfectly up to our max of 40 threads/node (Figure 1). Optimal scalability was calculated by projecting the walltime decrease proportionately to the increase in thread count, using the first walltime measurement at 4 threads as the starting point.

**Figure 1.**
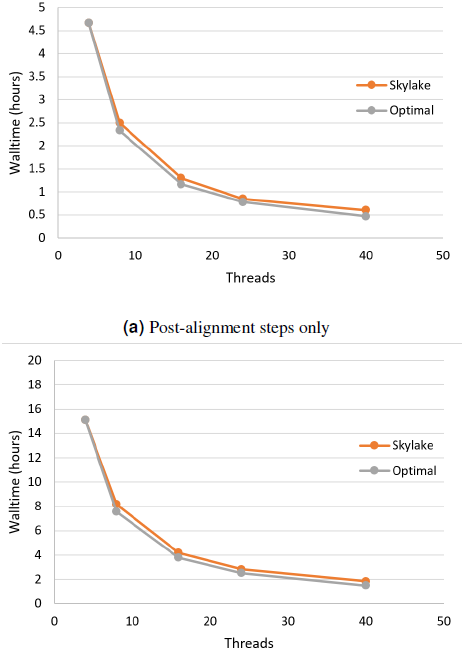
Sentieon DNASeq pipeline scales near-perfectly with the number of threads. Sample: NA12878, WGS, 20X. Datapoints reflect averages over two replicates.

### 2.5. Effect of sequencing depth on performance

Depth of sequencing coverage can have different impact at different steps in the Best Practices pipeline. Alignment proceeds in a linear fashion, one read at a time. Thus, higher coverage samples take a proportionately longer time to align. However, the local reassembly performed during realignment takes into account all reads at that locus simultaneously, which could result in a nonlinear relationship between walltime and coverage depth.

We investigated these relationships by running our Sentieon DNASeq pipeline on NA24694 WGS data subsets representing 25X, 50X, 75X, and 100X coverage depth. We ran each subset on the max available cores/node (40) to minimize the runtime. The individual tools in the pipeline again appeared to scale equally well (Figure 2a). The pipeline as a whole demonstrates near-linear scaling with depth. (Figure 2b).

**Figure 2.**
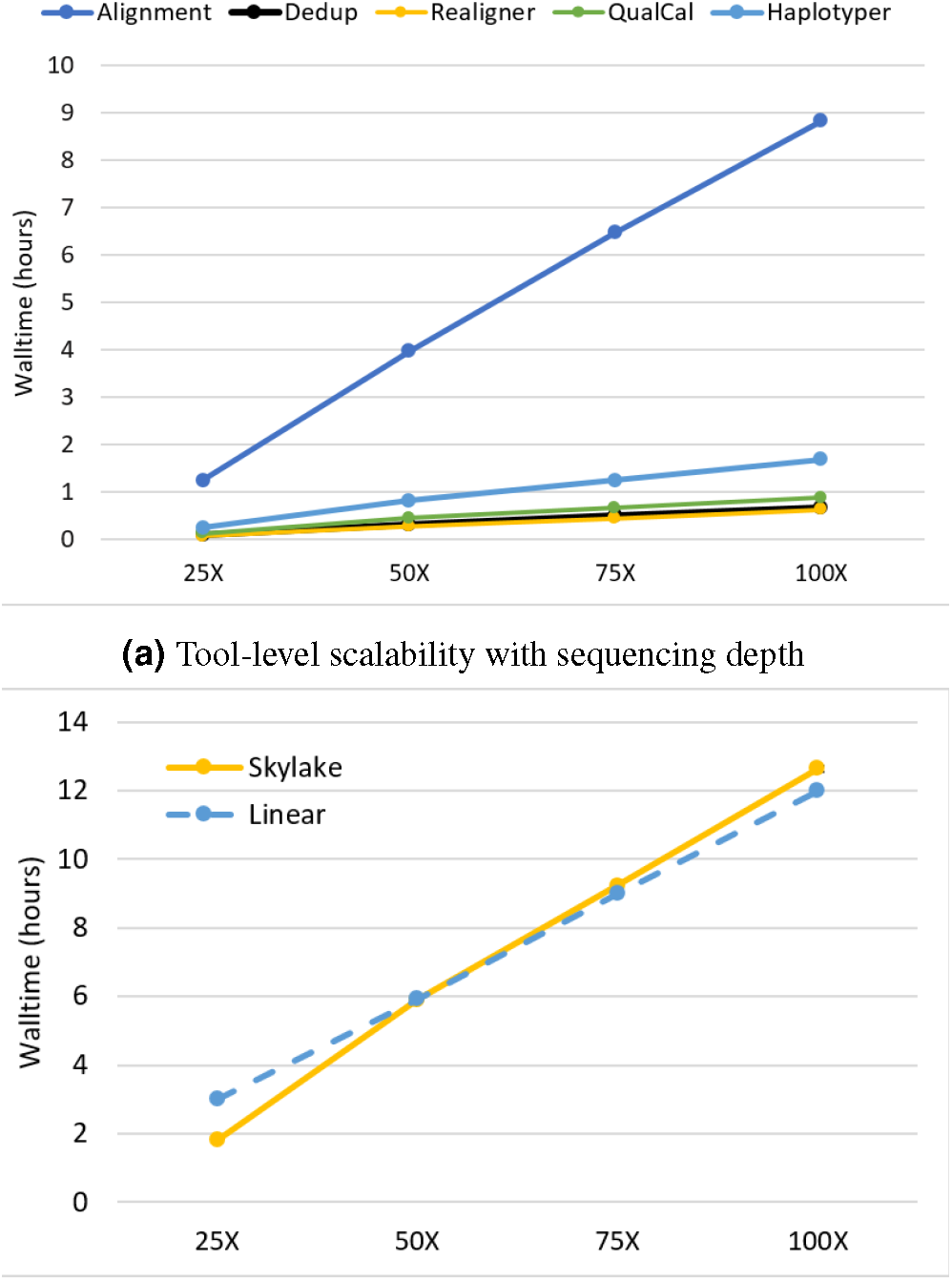
Sentieon DNASeq scalability as a function of sequencing coverage depth. Sample: NA24694, WGS, 25X-100X. Datapoints reflect averages over two replicates. Error bars are included in (b) but are too small to be visible.

### 2.6. Computational performance relative to GATK

To understand the extent of performance improvements introduced by Sentieon, we compared the runtime of GATK vs. DNASeq on NA12878 WGS data. The computational performance of GATK3.8 and GATK4 have been reviewed in detail by Heldenbrand et al. [6]. We ran each of the three pipelines with their respective default settings and maximum thread count (40) to establish a “baseline.” Then we ran each pipeline with “optimal” settings: for DNASeq, 40 threads across the pipeline, and for GATK3.8 and GATK4, the optimized parameters recommended in [6] (Table 3, reproduced with permission). No data-level parallelization was applied, and each test was performed on a single node.

Our walltime comparison excludes BWA MEM (not part of GATK; see section 2.7) and realignment (as the GATK team recommends against it). In this configuration, DNASeq runs for less than half an hour on NA12878 WGS 20X – an order of magnitude faster than GATK3.8 and GATK4, regardless of optimizations (Table 4). DNASeq runs in approximately 3% of the time taken by the fastest tested setup for GATK (GATK3.8 Optimized, 15.3h).

**Table 3.**
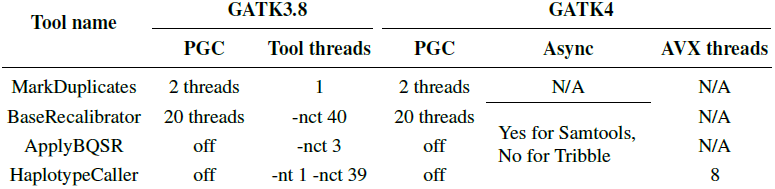
Summary of optimized parameter values for GATK3.8 and GATK4.

**Table 4.** Speed comparison: Sentieon DNASeq vs. GATK. Speedup factor is calculated on the DNASeq walltime relative to the corresponding GATK walltime.

| Pipeline | Walltime (hours) | Speedup |
| --- | --- | --- |
| DNaseSeq | .49 | - |
| GATK3.8 Baseline | 21.7 | x44 |
| GATK3.8 Optimized | 15.3 | x31 |
| GATK4.0 Baseline | 24.9 | x51 |
| GATK4.0 Optimized | 20.7 | x42 |

### 2.7. Comparison of Sentieon BWA to traditional BWA

Sentieon DNASeq includes an optimized version of BWA MEM, as well as a proprietary utility for sorting the aligned SAM into a BAM. This replaces an analogous pipeline starting with “traditional” BWA MEM [12] followed by samtools view [25], then NovoSort [26]. We compared performance of the two by running both with 40 threads on NA12878 WGS data (no piping). The Sentieon version was ∼28% faster: 1.25 hours vs. 1.73 hours. This speed-up results from optimizations to the klib library, at the cost of almost doubling the memory used by BWA: 22.45 ±1.58 GB vs 12.13 ±0.56 GB, measured as the resident set size (no swapping to disk was observed).

#### Version 201711.03

A new version of Sentieon (201711.03) was released as we completed our testing, featuring a nontrivial speed-up of BWA MEM derived from a complete rewrite of the traditional BWA MEM code. This new version runs 25% faster (0.95 hours) than 201711.02, and 45% faster than “BWA MEM → samtools view → NovoSort.”

### 2.8. Computer resource utilization

Variant calling workflows are notorious for having high RAM utilization, high rates of disk I/O, and inefficient data access patterns, which can cause tough performance issues [27, 28]. It is important to understand the patterns of compute resource utilization for any software attempting to perform variant calling. We recorded CPU load, memory utilization and I/O patterns for each tool while running the DNASeq pipeline on the NA12878 dataset. Our in-house profiling utility memprof [29] accesses /proc/PID for each process it monitors.

#### CPU utilization

The resultant profile (Figure 3) shows near-maximum core utilization by all tools except LocusCollector. This potentially indicates that the tools are largely CPU-bound, explaining the excellent thread-level scalability above.

**Figure 3.**
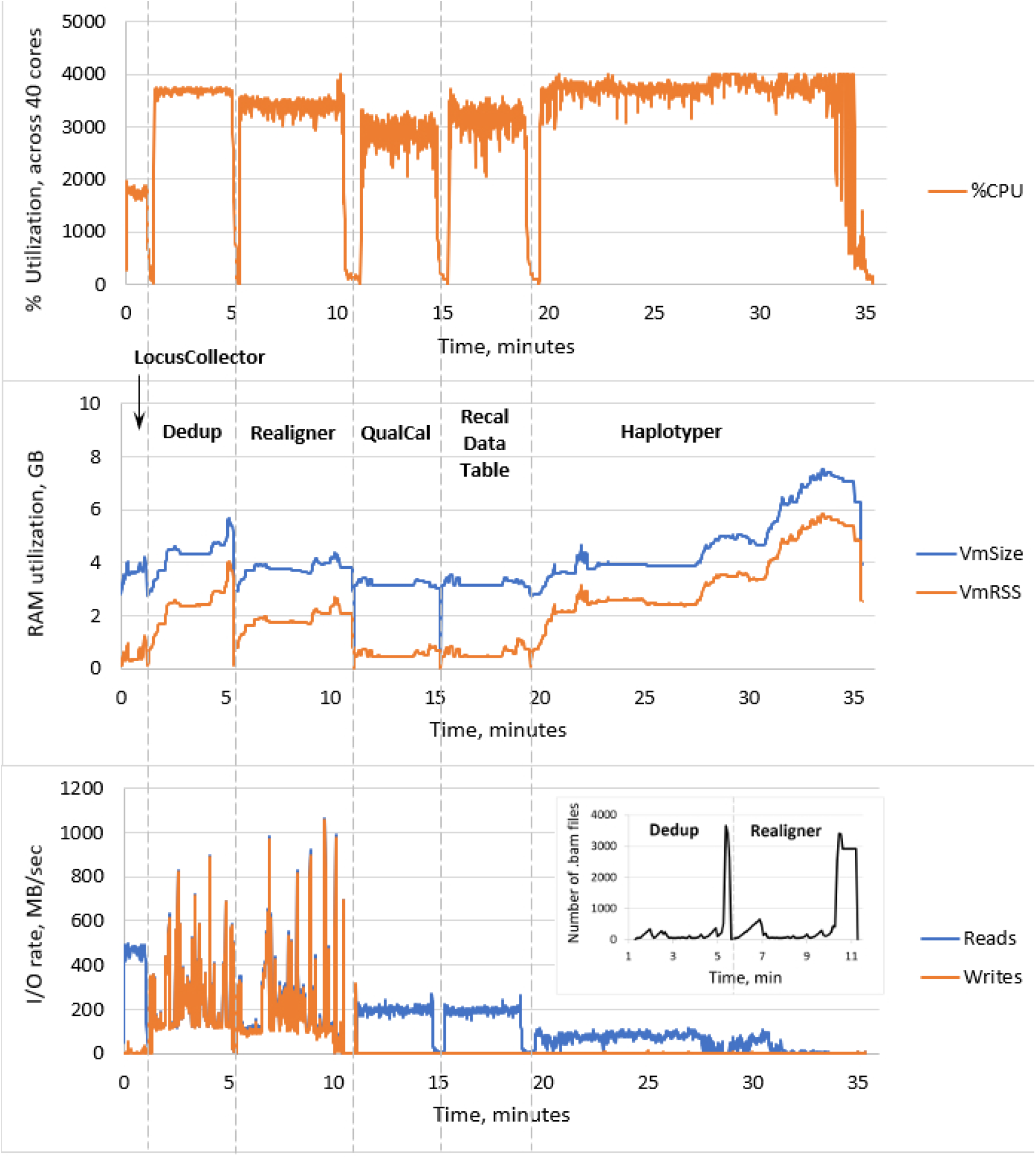
CPU utilization, memory usage and I/O of the Sentieon DNASeq tools, excluding BWA MEM. The pipeline steps are labeled in the middle panel, following the --algo option respectively used in the script. CPU utilization in the top panel corresponds to the sum total across the 40 cores on the node. RAM utilization in the middle panel was measured as resident set size (VmRSS) and total RAM reserved for computation (VmSize). I/O rates in the bottom panel were measures in reads and writes per second. Sample: NA12878, WGS, 20X.

#### RAM utilization

Haplotyper uses the most memory in the pipeline (up to 6 GB), as expected for the local reassembly subroutine. High RAM utilization toward the end of the process is likely due to processing of the difficult HG38 decoy regions. Nonetheless, DNASeq RAM utilization across all tools is lower than some previous GATK benchmarks [30].

#### I/O rates

Deduplication and realignment show very active I/O patterns, reaching high rates for both reads from the input BAM and writes to the output BAM. These patterns are extremely similar for the two steps (both for disk I/O and RAM I/O above), as they behave in similar ways: reading in a sequencing read, performing some simple operations and then writing a sequencing read to the disk. Many intermediary BAM files are created during these steps, resulting in high data-level parallelization at the cost of high I/O (Figure 3, inset in the bottom panel). This could introduce a filesystem bottleneck when analyzing large number of samples (hundreds) simultaneously on a cluster, but can be countered by using local disk or SSDs instead of network storage. In contrast, no new BAMs are created during QualCal, as it only calcu-lates the required modification of the quality scores, whereas the actual recalibration is applied during the variant calling stage. Thus the rate of writes/sec is very low for QualCal. For comparison we also included the near-identical profile for the optional command that applies the recalibration to calculate the post calibration data table (Figure 3, step **Recal Data Table**).

## 3. Conclusions

The tests presented here were intended to (a) benchmark Sentieon’s DNASeq speed and scalability and (b) compare DNASeq to GATK as an alternative option for variant calling. We determined that DNASeq scales optimally across threads, which makes it a powerful tool that will run well on a variety of processors. It also scales well, albeit suboptimally, as sequencing depth increases, a useful characteristic as deeper sequencing becomes more common. For a WGS sample sequenced to approximately 20X depth, DNASeq can complete the process from FASTQ to VCF in under 2 hours, and from aligned sorted BAM to VCF in less than half an hour. This opens up possibilities for point-of-care patient analysis in the clinic and massive reanalysis of legacy data.

Sentieon uses the same algorithms as GATK and reliably releases new versions in response to GATK version updates. Unlike GATK, which is open source, Sentieon software requires a license for use.

When compared to GATK, we found Sentieon DNASeq to be equally accurate. Comparisons to Illumina platinum calls for NA12878 yielded equivalent results, suggesting no meaningful differences in reliability. In terms of runtime, GATK post-alignment processing can take up to a day. DNASeq is able to complete the same work over 30x faster, representing a time savings of approximately 97%.

## Acknowledgments

This work was a product of the Mayo Clinic and Illinois Strategic Alliance for Technology-Based Healthcare. Major funding was provided by the Mayo Clinic Center for Individualized Medicine and the Todd and Karen Wanek Program for Hypoplastic Left Heart Syndrome. We thank the Interdisciplinary Health Sciences Institute, UIUC Institute for Genomic Biology and the National Center for Supercomputing Applications for their generous support and access to resources. We particularly acknowledge the support of Keith Stewart, M.B., Ch.B., Mayo Clinic/Illinois Grand Challenge Sponsor and Director of the Mayo Clinic Center for Individualized Medicine. Special gratitute to Amy Weckle for managing the project. Many thanks to the Sentieon team for the evaluation license, consultation and advice on the Sentieon variant calling software. The authors of this paper did not receive any benefits or compensation, financial or otherwise, from Sentieon in exchange for testing DNASeq, evaluating its performance, or expressing positive views thereof.

## References

[1] M. L. Metzker, “Sequencing technologies the next generation.” Nat Rev Genet, vol. 11, no. 1, pp. 31–46, Jan 2010. [Online]. Available: http://dx.doi.org/10.1038/nrg2626

[2] S. Goodwin, J. D. McPherson, and W. R. McCombie, “Coming of age: ten years of next-generation sequencing technologies.” Nat Rev Genet, vol. 17, no. 6, pp. 333–351, May 2016. [Online]. Available: http://dx.doi.org/10.1038/nrg.2016.49

[3] Z. D. Stephens, S. Y. Lee, F. Faghri, R. H. Campbell, C. Zhai, M. J. Efron, R. Iyer, M. C. Schatz, S. Sinha, and G. E. Robinson, “Big data: astronomical or genomical?” PLoS Biol, vol. 13, no. 7, p. e1002195, Jul 2015. [Online]. Available: http://dx.plos.org/10.1371/journal.pbio.1002195

[4] Illumina, “Illumina sequencing platforms,” 2018. [Online]. Available: https://www.illumina.com/systems/ sequencing-platforms.html

[5] The Broad Institute, “GATK | Best Practices,” 2017. [Online]. Available: https://software.broadinstitute.org/ gatk/best-practices/

[6] J. R. Heldenbrand, S. Baheti, M. A. Bockol, T. M. Drucker, S. N. Hart, M. E. Hudson, R. K. Iyer, M. T. Kalmbach, E. W. Klee, E. D. Wieben, M. Wiepert,D. E. Wildman, and L. S. Mainzer, “Performance benchmarking of GATK3.8 and GATK4,” bioRxiv, 2018. [Online]. Available: https://www.biorxiv.org/content/

[7] Sentieon DNASeq, “Sentieon,” 2018. [Online]. Available: https://www.sentieon.com/products/

[8] M. Plüss, A. M. Kopps, I. Keller, J. Meienberg, S. M. Caspar, N. Dubacher, R. Bruggmann, M. Vogel, and G. Matyas, “Need for speed in accurate whole-genome data analysis: GENALICE MAP challenges BWA/GATK more than PEMapper/PECaller and Isaac,” Proceedings of the National Academy of Sciences, p. 201713830, 2017.

[9] C. Raczy, R. Petrovski, C. T. Saunders, I. Chorny, S. Kruglyak, E. H. Margulies, H.-Y. Chuang, M. Källberg, S. A. Kumar, A. Liao, K. M. Little, M. P. Strömberg, and S. W. Tanner, “Isaac: ultra-fast whole-genome secondary analysis on Illumina sequencing platforms.” Bioinformatics, vol. 29, no. 16, pp. 2041–2043, aug 2013. [Online]. Available http://dx.doi.org/10.1093/bioinformatics/btt314

[10] M. A. DePristo, E. Banks, R. Poplin, K. V. Garimella, J. R. Maguire, C. Hartl, A. A. Philippakis, G. del Angel, M. A. Rivas, M. Hanna, A. McKenna, T. J. Fennell, A. M. Kernytsky, A. Y. Sivachenko, K. Cibulskis, S. B. Gabriel, D. Altshuler, and M. J. Daly, “A framework for variation discovery and genotyping using next-generation dna sequencing data.” Nat Genet, vol. 43, no. 5, pp. 491–498, May 2011. [Online]. Available: http://dx.doi.org/10.1038/ng.806

[11] G.A. Van der Auwera, M. O. Carneiro, C. Hartl, R. Poplin, G. Del Angel, A. Levy-Moonshine, T. Jordan, K. Shakir, D. Roazen, J. Thibault, E. Banks, K. V. Garimella, D. Altshuler, S. Gabriel, and M. A. DePristo, “From FastQ data to high confidence variant calls: the Genome Analysis Toolkit best practices pipeline,” Curr Protoc Bioinformatics, vol. 11, no. 1110, pp. 11.10.1–11.10.33, Oct 2013. [Online]. Available: http://dx.doi.org/10.1002/0471250953.bi1110s43

[12] H. Li, “Aligning sequence reads, clone sequences and assembly contigs with bwa-mem,” 2013. [Online]. Available: http://arxiv.org/abs/1303.3997v2

[13] “Broad institute’s software download page,” 2018. [Online]. Available: https://software.broadinstitute.org/ gatk/download/archive

[14] J. M. Zook, B. Chapman, J. Wang, D. Mittelman, O. Hofmann, W. Hide, and M. Salit, “Integrating human sequence data sets provides a resource of benchmark SNP and indel genotype calls,” Nature biotechnology, vol. 32, no. 3, p. 246, 2014.

[15] J. Zook, J. McDaniel, H. Parikh, H. Heaton, S. A. Irvine,L. Trigg, R. Truty, C. Y. McLean, F. M. De La Vega, C. Xiao, S. Sherry, and M. Salit, “Reproducible integration of multiple sequencing datasets to form high-confidence SNP, indel, and reference calls for five human genome reference materials,” bioRxiv, 2018. [Online]. Available:https://www.biorxiv.org/content/early/2018/05/25/281006

[16] G. M. Church, “The personal genome project,” Molecular systems biology, vol. 1, no. 1, 2005.

[17] “Genome in a bottle—a human dna standard,” Nature biotechnology, vol. 33, p. 675, 2015. [Online]. Available: http://www.nature.com.proxy2.library.illinois. edu/articles/nbt0715-675a

[18] Z. D. Stephens, M. E. Hudson, L. S. Mainzer, M. Taschuk, M. R. Weber, and R. K. Iyer, “Simulating next-generation sequencing datasets from empirical mutation and sequencing models,” PLOS ONE, vol. 11, no. 11, pp. 1–18, Nov 2016. [Online]. Available: https://doi.org/10.1371/journal.pone.0167047

[19] Z. Stephens, “neat-genreads,” 2018. [Online]. Available: https://github.com/zstephens/neat-genreads

[20] “PrecisionFDA Truth Challenge,” 2016. [Online]. Available: https://precision.fda.gov/challenges/truth/ results

[21] “PrecisionFDA Consistency Challenge,” 2016. [Online]. Available: https://precision.fda.gov/challenges/ consistency/results

[22] S. Bionetworks, “ICGC-TCGA DREAM Mutation Calling Challenge,” 2016. [Online]. Available: https://www.synapse.org/#!Synapse:syn312572/wiki/247695

[23] Z. Stephens, “NEAT vcf-compare,” 2015. [Online]. Available: https://web.engr.illinois.edu/∼zstephe2/read_simulator/vcfCompare.html

[24] “Illumina platinum confident regions,” 2018. [Online]. Available: https://github.com/Illumina/ PlatinumGenomes/blob/master/files/2017-1.0.files

[25] H. Li, B. Handsaker, A. Wysoker, T. Fennell, J. Ruan, N. Homer, G. Marth, G. Abecasis, R. Durbin, and. G. P. D. P. Subgroup, “The sequence alignment/map format and samtools.” Bioinformatics, vol. 25, no. 16, pp. 2078–2079, Aug 2009. [Online]. Available: http://dx.doi.org/10.1093/bioinformatics/btp352

[26] NOVOCRAFT TECHNOLOGIES SDN BHD, “Novocraft,” 2014. [Online]. Available: http://www.novocraft.com/

[27] S. S. Banerjee, A. P. Athreya, L. S. Mainzer, C. V. Jongeneel, W.-M. Hwu, Z. T. Kalbarczyk, and R. K. Iyer, “Efficient and scalable workflows for genomic analyses,” in Proceedings of the ACM International Workshop on Data-Intensive Distributed Computing. ACM, 2016, pp. 27–36.

[28] N. Kathiresan, R. Temanni, H. Almabrazi, N. Syed, P. V. Jithesh, and R. Al-Ali, “Accelerating next generation sequencing data analysis with system level optimizations,” Scientific Reports, vol. 7, no. 1, p. 9058, 2017.

[29] V. Kindratenko, “Performance profiling utility memprof,” 2018. [Online]. Available:https://github.com/IGBIllinois/memprof

[30] Brad Chapman, “Benchmarking variation and rna-seq analyses on amazon web services with docker,” 2014. [Online]. Available:https://www.sentieon.com/products/

